# NMDAR-activated PP1 dephosphorylates GluN2B to modulate NMDAR synaptic content

**DOI:** 10.1101/520882

**Authors:** Andrew M. Chiu, Jiejie Wang, Michael P. Fiske, Pavla Hubalkova, Levi Barse, John A. Gray, Antonio Sanz-Clemente

## Abstract

In mature neurons, postsynaptic NMDARs are segregated into two populations, synaptic and extrasynaptic, which differ in localization, function, and associated intracellular cascades. These two pools are connected via lateral diffusion, and receptor exchange between them modulates synaptic NMDAR content. Here, we identify the phosphorylation of the PDZ-ligand of the GluN2B subunit of NMDARs (at S1480) as a critical determinant in dynamically controlling NMDAR synaptic content. We find that phosphorylation of GluN2B at S1480 maintains NMDARs at extrasynaptic membranes as part of a protein complex containing Protein Phosphatase 1 (PP1). Global activation of NMDARs leads to the activation of PP1, which mediates dephosphorylation of GluN2B at S1480 to promote an increase in synaptic NMDAR content. Thus, PP1-mediated dephosphorylation of the GluN2B PDZ-ligand modulates the synaptic expression of NMDARs in mature neurons in an activity-dependent manner, a process with profound consequences for synaptic and structural plasticity, metaplasticity, and synaptic neurotransmission.

**HIGHLIGHTS:**

- Phosphorylation of the PDZ-ligand of the GluN2B subunit of NMDARs (GluN2B-pS1480) maintains NMDARs at extrasynaptic sites.
- Extrasynaptic NMDARs form a stable protein complex containing PP1.
- Global NMDAR activation increases NMDAR synaptic content by promoting PP1-mediated dephosphorylation of GluN2B-pS1480.
- GluN2B-pS1480 dephosphorylation is mediated by a PP1 subpopulation not involved in LTD and is enhanced by age.

## INTRODUCTION

N-methyl-D-aspartate receptors (NMDARs) are ionotropic glutamate receptors essential for excitatory neurotransmission in the brain (Paoletti et al., 2013). In mature neurons, functional postsynaptic NMDARs are segregated in two different locations: synaptic NMDARs, which are encapsulated within the postsynaptic density (PSD), and extrasynaptic NMDARs, which are distributed between perisynaptic sites and dendritic shafts (Papouin and Oliet, 2014). These two populations have distinct functions and trigger different intracellular cascades. For example, synaptic NMDAR activation induces classical forms of synaptic plasticity (i.e., LTP and LTD) and activates ERK and CREB signaling, whereas extrasynaptic NMDARs have neuronal modulatory roles and inhibit ERK and CREB pathways (Parsons and Raymond, 2014). The NMDAR content in both populations is plastic and can be dynamically modified in response to different stimuli (Paoletti et al., 2013). The paradigmatic example is the switch in the synaptic GluN2 subunit composition of NMDARs that occurs at early developmental stages (from predominantly GluN2B-containing to predominantly GluN2A-containing NMDARs) which decisively contributes to synaptic maturation (Sanz-Clemente et al., 2013b). Growing evidence shows that synaptic NMDARs are subject to activity-dependent regulation in mature neurons as well (Hunt and Castillo, 2012). This bidirectional plasticity of NMDARs has been hypothesized to affect important synaptic functions such as metaplasticity and homeostatic plasticity, and to modify the integrative properties of the neuron (Dore et al., 2017; Hunt and Castillo, 2012). Mechanistically, both an increase in surface-expressed NMDARs and the synaptic recruitment of extrasynaptic NMDARs contributes to an increase of NMDAR synaptic content (Harney et al., 2008; Tovar and Westbrook, 2002). However, the molecular determinants controlling these processes are not fully understood.

We have previously identified a molecular mechanism controlling the clearance of GluN2B-containing NMDARs from synaptic sites by showing that the phosphorylation of the GluN2B PDZ-ligand (at S1480; GluN2B-pS1480) promotes the lateral diffusion of synaptic NMDARs to extrasynaptic sites (Chen et al., 2012; Sanz-Clemente et al., 2013a; Sanz-Clemente et al., 2010). We report here that GluN2B-pS1480-containing NMDARs are expressed at extrasynaptic membranes and they must be dephosphorylated in order to enter into synaptic sites. Additionally, we have identified Protein Phosphatase 1 (PP1) as the phosphatase mediating this reaction and outlined the molecular determinants that control this process by showing that extrasynaptic NMDARs form a stable protein complex containing PP1. Global activation of NMDARs induces dephosphorylation of GluN2B-pS1480 by activating a subpopulation of PP1 which is not involved in LTD and promotes the insertion of extrasynaptic GluN2B receptors into the PSD. This novel mechanism, therefore, controls the activity-dependent NMDAR-plasticity in mature neurons.

## RESULTS

We have previously identified a molecular mechanism promoting the clearance of GluN2B-containing NMDARs from PSDs, which is based on the synaptic activity-dependent phosphorylation of the GluN2B subunit within its PDZ-ligand domain (at S1480; GluN2B-pS1480) mediated by casein kinase 2 (CK2). Briefly, GluN2B-pS1480 disrupts the interaction of NMDARs with synaptic scaffolding proteins (e.g., PSD-95), which facilitates lateral diffusion to extrasynaptic sites, where the phosphorylated receptors are internalized in a clathrin-dependent manner (Chen et al., 2012; Sanz-Clemente et al., 2013a; Sanz-Clemente et al., 2010). Because GluN2B-containing NMDARs are fated for recycling (Lavezzari et al., 2004) and GluN2B-pS1480 seems to be permissive for lateral diffusion between synaptic and extrasynaptic populations, we initially hypothesized that internalized NMDARs would be recycled and reinserted into the PSD. This idea would be consistent with the demonstrated GluN2B-containing NMDARs exchange between synaptic and extrasynaptic populations via lateral diffusion (Dupuis et al., 2014).

We first tested this hypothesis by using an antibody feeding approach in primary hippocampal neurons overexpressing GFP-GluN2B wild-type (WT) or the phospho-mimetic mutant GluN2B S1480E, which, despite comparable total expression to GluN2B WT, displays impaired surface expression due to increased internalization (Sanz-Clemente et al., 2010). Consistent with our hypothesis, we found that the phospho-mimetic mutant was readily recycled to the plasma membrane to a level comparable to GluN2B WT (Figure 1A). Next, as a first step to test the reinsertion of phosphorylated receptors into the PSD, we evaluated NMDAR surface and synaptic expression by conducting genetic molecular replacement experiments in floxed GluN2A/2B mice as previously described. (Chen et al., 2012). Briefly, we used organotypic hippocampal slices biolistically co-transfected with either Cre alone, to remove endogenous GluN2A and GluN2B, or Cre with exogenous GluN2B S1480E. Then, we measured both evoked NMDAR-mediated synaptic currents (NMDAR-EPSCs) and whole cell responses to puffed-on NMDA. As we have previously shown (Chen et al., 2012), we found that neurons expressing only the phospho-mimetic mutant GluN2B S1480E largely lacked NMDAR-EPSCs (Figure 1B), but we found that whole cell responses to puffed-on NMDA were present in neurons expressing GluN2B S1480E (Figure 1C) demonstrating surface expression of GluN2B S1480E.

**FIGURE 1:**
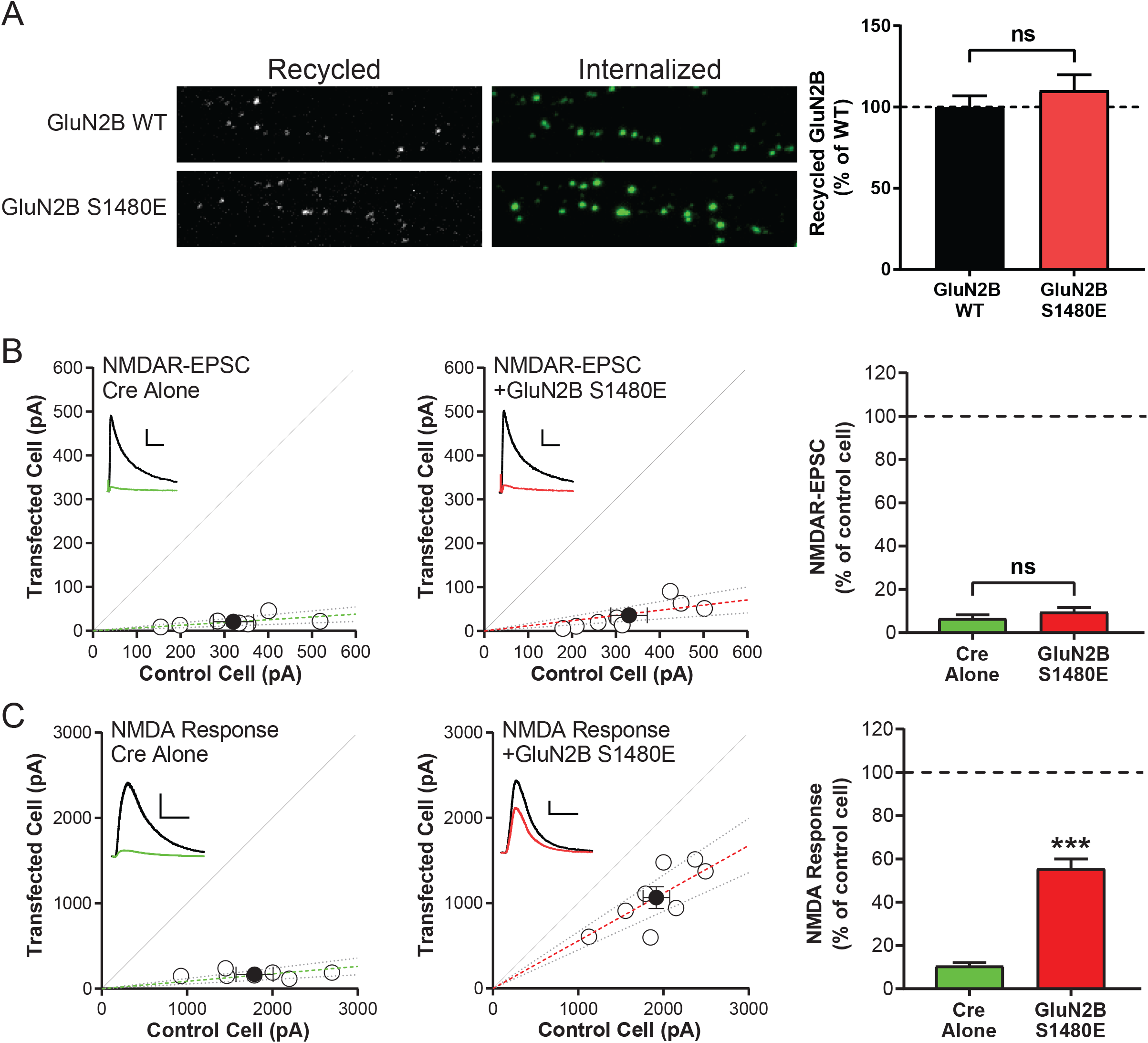
Phosphorylation of the PDZ-ligand of the GluN2B subunit (GluN2B-pS1480) of NMDARs maintains NMDARs segregated at extrasynaptic membranes. All graphs represent mean ± SEM. ns=non-significant, ***p<.001 in a Mann-Whitney U test. **A)** Hippocampal neurons were transfected at DIV7 with GFP-GluN2B WT or the phospho-mimetic mutant S1480E. At DIV13-14, the recycling ratio (recycled/internalized) of the exogenous receptors was analyzed using an antibody-feeding approach (see Methods for details). N=3, n (WT, S1408E)=22, 20. p=.54 **B)** Organotypic hippocampal slices were prepared from P7 *Grin2a*^fl/fl^*Grin2b*^fl/fl^ mice, biolistically transfected with either Cre or Cre + GluN2B-S1480E at DIV2-4, and paired whole-cell recordings were obtained from Cre-expressing and neighboring control CA1 pyramidal neurons at DIV18-24. Scatter plots represent peak amplitudes of NMDAR-EPSCs from single pairs (open circles) and mean ± SEM (filled circles) from transfected and control cells. Dashed lines represent linear regression and 95% confidence interval. For sample traces, control cells are represented black and transfected cells are represented in green/red; scale bars represent 100 ms and 100 pA. p=0.54. **C)** After NMDAR-EPSC measurement, whole-cell responses to 100 µM NMDA puffed onto the same cells as in (B). Scatter plots represent peak amplitudes of NMDA responses from single pairs (open circles) and mean ± SEM (filled circles) from transfected and control cells. Dashed lines represent linear regression and 95% confidence interval. Sample traces are as in (B); scale bars represent 5 sec and 400 pA.

These data indicate that, while a population of phosphorylated receptors can be trafficked to the surface, these receptors are not stabilized at PSDs and are exclusively present at extrasynaptic sites. Based on these data, we formulated our central hypothesis: surface expressed GluN2B-containing NMDARs must be dephosphorylated within the GluN2B PDZ-ligand prior to integration into the PSD.

### Global activation of NMDARs promotes Protein Phosphatase 1 (PP1)-mediated dephosphorylation of S1480 on surface-expressed GluN2B subunits

If GluN2B needs to be dephosphorylated to re-enter PSDs, then this reaction should occur on surface-expressed GluN2B-containing NMDARs. To test our hypothesis, we first induced GluN2B dephosphorylation in primary cortical cultures by globally activating NMDARs as previously described (bath application of 50 µM NMDA for 10 min) (Chung et al., 2004). As expected, acute NMDA treatment did not affect the total expression of GluN2B (not shown), but it promoted a robust dephosphorylation of GluN2B-pS1480 in whole cell lysate (Figure 2). Next, we isolated surface-expressed NMDARs using two independent methods: biotinylation (Figure 2A) and biochemical isolation of the synaptic plasma membranes (SPMs) (Figure 2B). Finally, we quantified the levels of GluN2B-pS1480 by immunoblotting using a custom-generated antibody against the phosphorylated state of GluN2B at S1480 (Figure S1). Importantly, the reduction in GluN2B phosphorylation was similar in total lysate and in surface-expressed proteins, indicating that global NMDAR activation dephosphorylates the NMDAR population with the potential to be integrated into the PSD through lateral insertion.

**FIGURE 2:**
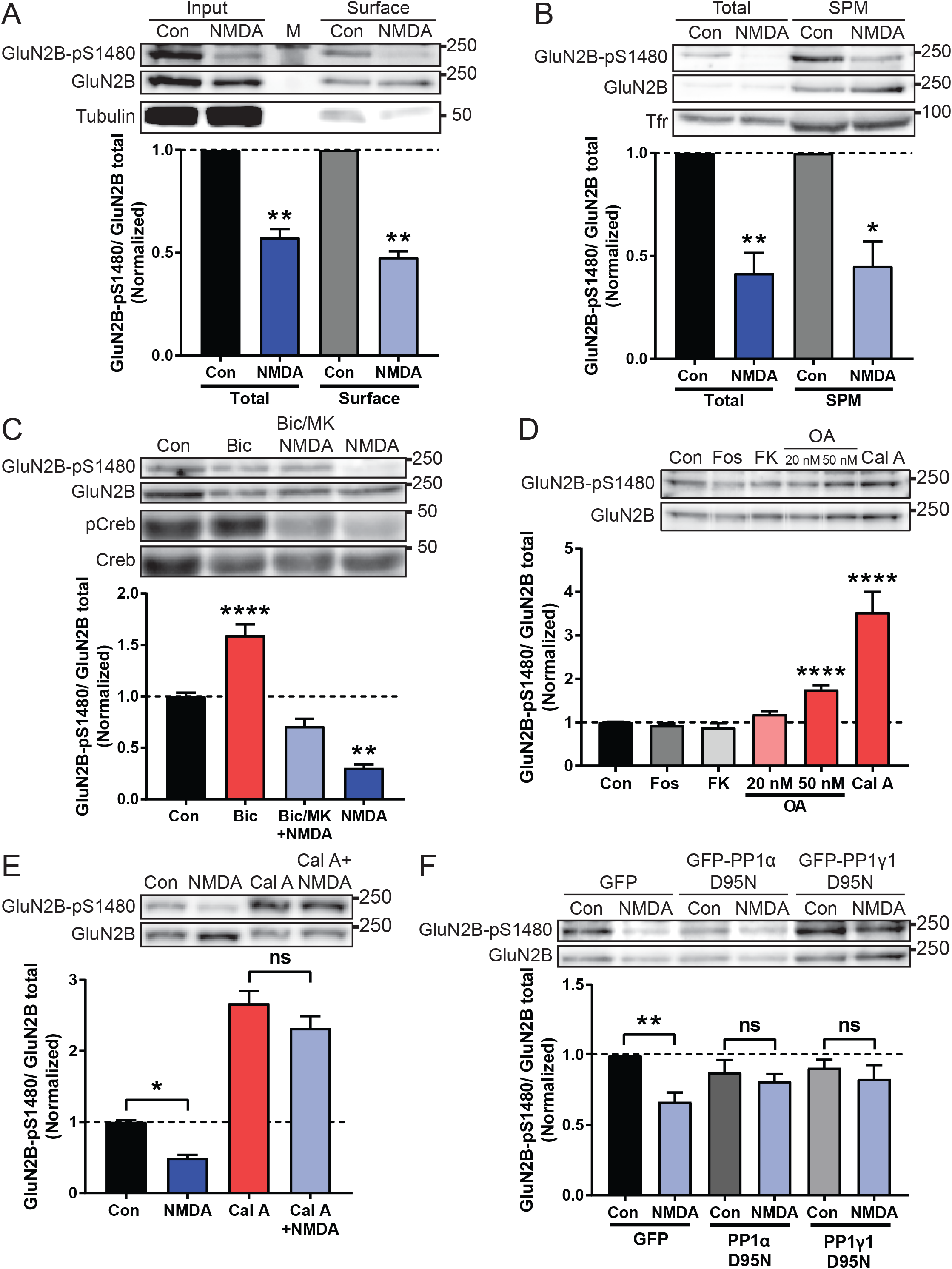
Co-activation of synaptic and extrasynaptic NMDARs promotes Protein Phosphatase 1 (PP1)-mediated dephosphorylation of S1480 on surface-expressed GluN2B. Levels of normalized GluN2B-pS1480 were analyzed by immunoblotting from cortical primary neurons after the indicated manipulations. Graphs represent mean ± SEM. *p<.05, **p<.01, ****p<.0001 using a Mann-Whitney U test (A and B) or Kruskal-Wallis H test (C-F). **A)** Biotinylation experiment in DIV24-28 cortical neurons after induction of GluN2B-pS1480 dephosphorylation by NMDA incubation (50 µM for 10 min). Tubulin shown as a control. M denotes marker. N=5 **B)** Isolation of the synaptic plasma membrane (SPM) fraction from DIV14-21 cortical neurons following NMDA treatment as before. Transferrin receptor (Tfr) shown as a control. N=5 **C)** Synaptic (50 µM Bicuculline for 20 min), Extrasynaptic (50 µM NMDA for 10 min after blocking synaptic NMDARs by incubation with 50 µM Bicuculline in the presence of 10 µM MK-801 for 10 min) or Global (50 µM NMDA for 10 min) activation of NMDARs in DIV21-28 cortical neurons. pCreb and Creb shown as controls. N=3 **D)** Incubation of DIV14-17 neurons with phosphatase inhibitors for 45 min: Fostriecin for PP2A/4 (Fos, 1 µM), FK506 for PP2B (FK, 1 µM), Okadaic Acid for PP2A at 20 nM and PP2A/PP1 at 50 nM (OA, 20 or 50 nM), and Calyculin A for PP1/PP2A (Cal A, 100 nM). N=10 **E)** Preincubation of DIV14-21 cortical neurons with 100 nM Cal A for 45 min before induction of GluN2B-pS1480 dephosphorylation by bath application of NMDA as before. p=.66 (Cal A vs Cal A + NMDA). N=14 **F)** Lentiviral transduction of the dominant negative forms of GFP-PP1 α and γ1 (PP1α/γ1 D95N) and GFP (as a control) in DIV14-17 cortical neurons for 7-10 days. GluN2B-pS1480 dephosphorylation was induced with NMDA as before. p>.99 (PP1α) and p=.78 (PP1γ1). N=6 Related to Figure S1 and S2

Because previous studies have shown that synaptic NMDAR activation induces GluN2B S1480 phosphorylation and global NMDAR activation promotes dephosphorylation (Chung et al., 2004; Sanz-Clemente et al., 2010) (Figures 2A/B), we wondered if selective activation of extrasynaptic NMDARs would be sufficient for GluN2B-pS1480 dephosphorylation. Thus, we utilized an established protocol for the preferential stimulation of extrasynaptic NMDARs in primary cortical cultures and quantified phosphorylated CREB levels as a control (Hardingham et al., 2002). Briefly, we inhibited synaptic NMDARs by promoting their opening in the presence of the irreversible NMDAR blocker MK-801. After MK-801 withdrawal, remaining functional NMDARs (i.e., mostly extrasynaptic) were activated by bath application of NMDA as before. This protocol was unable to maximally dephosphorylate GluN2B-pS1480 (Figures 2C), suggesting that both synaptic and extrasynaptic NMDAR populations synergize to dephosphorylate GluN2B. Because a major consequence of GluN2B-pS1480 is the decrease in GluN2B surface expression due to an enhanced internalization ratio (Sanz-Clemente et al., 2010), we next investigated if the pharmacological manipulations that control GluN2B-pS1480 also affect GluN2B surface expression. Consistent with our biochemical data (Figure 2C), global activation of NMDARs by NMDA treatment resulted in a robust increase in surface-expressed GluN2B, whereas preferential activation of extrasynaptic NMDARs failed to significantly modify GluN2B surface expression (Figure S2).

We finally sought to identify the phosphatase directly responsible for GluN2B-pS1480 dephosphorylation. Because the majority of dephosphorylation reactions are controlled by the phosphoprotein phosphatase (PP) family (Virshup and Shenolikar, 2009), we first used a pharmacologic approach to inhibit PPs in neuronal cultures and probed for changes in baseline GluN2B-pS1480. As shown in Figure 2D, we observed a robust increase in GluN2B-pS1480 in neurons treated with okadaic acid (OA, 50 nM) and calyculin A (Cal A, 100 nM). At the concentrations used, both these inhibitors block protein phosphatase 1 (PP1) and protein phosphatase 2A (PP2A). Because treatment with fostriecin, a PP2A and PP4 inhibitor, was unable to cause changes in GluN2B S1480 phosphorylation, we hypothesized that PP1 is the phosphatase responsible for GluN2B-pS1480 dephosphorylation. Supporting this idea, low concentrations of OA (under 50 nM) which preferentially inhibit PP2A (Ishihara et al., 1989), did not modulate GluN2B-pS1480 (Figure 2D). Importantly, pre-incubation with Cal A (Figure 2E) or 50 nM OA (not shown) blocked the NMDA-triggered dephosphorylation of GluN2B-pS1480, demonstrating that PP1 also controls the agonist-driven reaction. We confirmed the involvement of PP1 in GluN2B-pS1480 dephosphorylation by using a genetic approach. Specifically, we transduced cultured neurons with lentiviruses expressing the dominant negative form of the two PP1 isoforms present in dendrites [PP1α and PP1γ1 (D95N)] (Strack et al., 1999; Zhang et al., 1996) and tested the levels of GluN2B-pS1480 by immunoblotting. Overexpression of nonfunctional PP1 prevented NMDA-driven dephosphorylation of GluN2B S1480, supporting that PP1 is the phosphatase responsible for GluN2B-pS1480 dephosphorylation (Figure 2F).

Together, our data demonstrate that global activation of NMDARs triggers the PP1-mediated dephosphorylation of GluN2B-pS1480 on NMDARs expressed at the cell surface.

### Regulation of the PP1-dependent dephosphorylation of GluN2B S1480

PP1 is a constitutively active phosphatase with little inherent specificity. Thus, the cell utilizes several molecular mechanisms to regulate PP1 activity (Figure 3A) (Cohen, 2002). These include (1) direct inhibitory phosphorylation of the PP1 c-terminus, a reaction canonically controlled by CDK5, (2) complexing with inhibitor proteins to block accessibility to the active site, and (3) complexing of active PP1 with target specifying proteins to confer specificity and restrict PP1 to a specific subcellular compartment. We next sought to examine which of these modes of PP1 regulation participate in GluN2B-pS1480 dephosphorylation.

**FIGURE 3:**
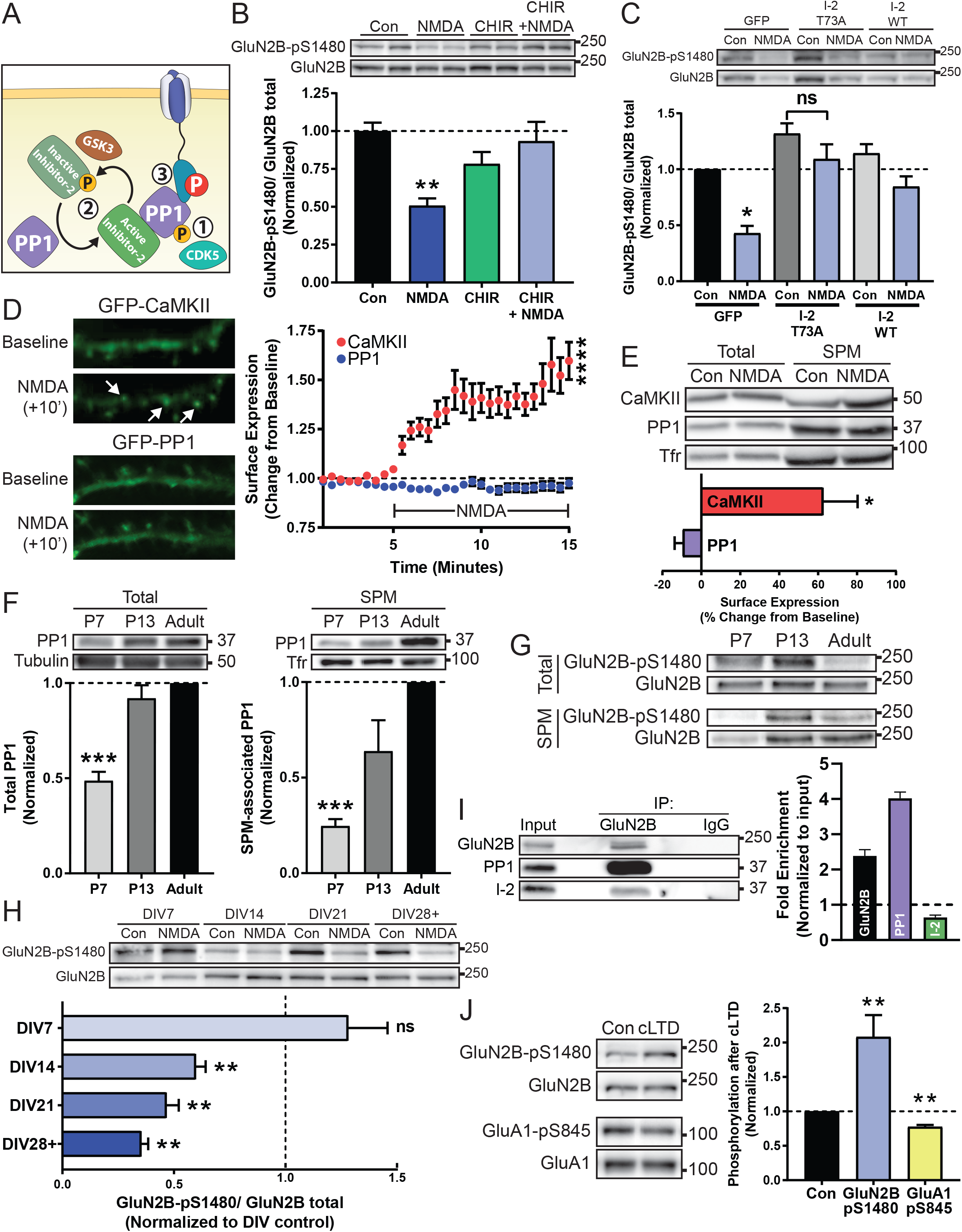
Regulation of the PP1-dependent dephosphorylation of GluN2B S1480. All graphs represent mean ± SEM. ns=non-significant, *p<.05, **p<.01, ***p<.001, ****p<.0001 in a Kruskal-Wallis H test (B and G) or a Mann-Whitney U test (C, D, and F). **A)** PP1 is regulated by (1) inhibitory CDK5 phosphorylation (2) complexing with endogenous inhibitor proteins or (3) targeting to a correct substrate. **B)** Levels of normalized GluN2B-pS1480 were analyzed by immunoblotting from DIV14 cortical primary neurons after pharmacological blockade of Inhibitor-2 (I-2) inactivation by blocking GSK3 (CHIR99021, 50 µM for 2 hours) prior to NMDA-triggered dephosphorylation of GluN2B-pS1480. N=8 **C)** Levels of normalized GluN2B-pS1480 were analyzed by immunoblotting from DIV14-17 cortical primary neurons after lentiviral transduction of the constitutively active form of GFP-I-2 (I-2 T73A) or GFP and GFP-I-2 WT (as controls) in cortical neurons for 7-10 days. GluN2B-pS1480 dephosphorylation was induced with NMDA as before. p=.63 (I-2 T73A). N=6 **D)** NMDA-induced translocation to the plasma membrane of GFP-PP1 and GFP-CaMKII (as a control) was evaluated in DIV14 hippocampal cultures by Total Internal Reflection Fluorescence (TIRF) microscopy. Baseline fluorescence was recorded for 5 mins prior to the addition of 50 μM NMDA as before. Graph represents changes in TIRF signal over time. N=4, n=11. Related to Movies S1 and S2. **E)** The synaptic plasma membrane (SPM) fraction of DIV14-21 cortical neurons was isolated and the relative expression of the indicated proteins was evaluated by immunoblotting. Protein levels were normalized to transferrin receptor (Tfr) in each condition and then normalized to control surface expression levels. N=4 **F)** Levels of PP1 present in the SPM fraction of mice of the indicated age were assessed by immunoblotting. N=8 **G)** Levels of GluN2B-pS1480 present in the SPM fraction of mice of the indicated age were assessed by immunoblotting. Graph represents fold of enrichment normalized to “input”. N=3 **H)** Levels of normalized GluN2B-pS1480 were analyzed by immunoblotting from cortical primary neurons after GluN2B-pS1480 dephosphorylation was induced in cortical neurons of the indicated DIV by incubating with 50 µM NMDA for 10 min as before. p=.68 (DIV7). N=5 **I)** The extrasynaptic fraction from rat cortex was isolated and the association of GluN2B, PP1 and I-2 was analyzed by immunoblotting following co-immunoprecipitation with an anti-GluN2B antibody. **J)** cLTD (20 μM NMDA for 3 mins and 30 min recovery) was initiated in acute hippocampal slices from P72-P84 mice. Levels of GluN2B-pS1480 and GluA1-pS845 were assessed by immunoblotting. N=6. Related to Figure S3

CDK5-mediated phosphorylation of PP1 (at T320) inhibits phosphatase activity by blocking the access of PP1 to its substrates. As previously reported (Hou et al., 2013), we found that bath application of NMDA to primary neuronal cultures causes a global reduction in PP1 T320 phosphorylation (i.e., increasing the amount of active PP1) (Figure S3A).

Neurons express several inhibitor proteins which complex with PP1 to block access to the active site, including inhibitor 1 (I-1), inhibitor-2 (I-2), and dopamine- and cAMP-regulated phosphoprotein-32 (DARPP-32). In turn, these inhibitors are regulated by kinase and phosphatase activity to allow for signal integration (Eto and Brautigan, 2012). The involvement of I-1 and DARPP-32 in GluN2B-pS1480 regulation is unlikely because of their relatively low expression in hippocampal CA1 neurons (Glausier et al., 2010). Nevertheless, pharmacological manipulations in culture to modulate I-1 and DARPP-32 activation failed to control GluN2B-pS1480 dephosphorylation, excluding them as molecular players in this process (Figures S2B-D). Next, we tested the involvement of Inhibitor 2 (I-2), which functions as an active inhibitor in its dephosphorylated state (Siddoway et al., 2013). GSK3β-mediated phosphorylation of I-2 inactivates the inhibitor and it is known that global NMDAR activation causes both an increase in active GSK3β and an increase in phosphorylated I-2 (Szatmari et al., 2005). I-2, therefore, is an appealing candidate for PP1 inhibition in our reaction. As before, we first used a pharmacological approach and inhibited GSK3β in neuronal cultures using the GSK3 specific inhibitor CHIR99021 (10 μM for 2 hours). GSK3 blockade prevented the NMDA-mediated GluN2B-pS1480 dephosphorylation, suggesting a role for I-2 in this process (Figures 3B). To further confirm I-2 involvement, we generated lentiviruses expressing a phospho-deficient form of I-2 (I-2 T73A) to inhibit PP1 constitutively. As expected, overexpression of this mutant prevented NMDA-triggered GluN2B-pS1480 dephosphorylation (Figures 3C).

The last key piece of PP1 regulation is correct targeting and complexing of PP1 with its substrate (Hendrickx et al., 2009). Because PP1 is canonically thought of as a cytosolic protein which is not associated to the plasma membrane but dephosphorylation occurs on NMDARs expressed at the cell surface (Figure 2A/B), we hypothesized that PP1 might translocate to a plasma membrane-associated population in response to NMDAR activation. To examine this possibility, we first utilized Total Internal Reflection Fluorescence (TIRF) microscopy in primary hippocampal cultures. This imaging technique enables examination of dynamic processes that occur within 100 nm of the plasma membrane. We overexpressed GFP-PP1 in hippocampal neurons and tracked its targeting to the surface in response to NMDA treatment over time. As a control, we used GFP-CaMKII, as the translocation of this kinase to the synapses after NMDAR activation has been extensively characterized (Merrill et al., 2005). As expected, CaMKII showed robust surface trafficking in response to NMDA treatment, but surprisingly, PP1 showed no change in surface levels (Figure 3D). Because overexpression of GFP-PP1 may saturate the system, we next isolated the SPM fraction following NMDA treatment in primary cortical neurons and quantified the levels of endogenous PP1 associated with SPMs by immunoblotting (Figures 3E). Again, we observed a strong enrichment of CaMKII in the SPM fraction after NMDA treatment, with no increase in SPM-associated PP1. These data indicate that the PP1 population responsible for GluN2B S1480 dephosphorylation is present at the plasma membrane, likely, already associated with NMDAR-complexes.

### GluN2B-pS1480 dephosphorylation is enhanced by neuronal maturation

Our finding that GluN2B-pS1480 dephosphorylation is dependent on PP1 associated with SPMs resembles the regulation of the complementary process, phosphorylation of GluN2B at S1480, which we have previously shown to be strongly regulated by the level of SPM-associated CK2 (Sanz-Clemente et al., 2013a). Interestingly, we have previously demonstrated that the association of CK2 with SPMs is developmentally regulated and it peaks around the second postnatal week in rodents, coincident with the highest level of GluN2B-pS1480 and the developmental switch in synaptic GluN2 subunit composition (Sanz-Clemente et al., 2010). Therefore, we next investigated if the association of PP1 with SPMs also has a developmental component and found that low amounts of PP1 were detectable in the SPM fraction isolated from immature mouse brain (P7), whereas the association of PP1 with SPMs dramatically increased throughout brain maturation. This is not a mere consequence of changes in PP1 total expression, as PP1 is strongly expressed since early development (Figure 3F). These data suggest that the developmentally controlled association to SPMs of both the kinase (CK2) and the phosphatase (PP1) synergize to reduce the levels of GluN2B-pS1480 during the adulthood (Figure 3G) (Sanz-Clemente et al., 2010).

The increasing level of SPM-associated PP1 throughout development suggests that, in addition to the developmental changes in basal GluN2B-pS1480, the NMDAR activity-driven GluN2B-pS1480 dephosphorylation might be differentially controlled at different stages of synaptic maturation. We tested this possibility by inducing NMDA-dependent GluN2B-pS1480 dephosphorylation in primary cortical neurons at different stages of maturation (from DIV7 to DIV28-29). As expected, we found a strong influence of the maturation stage in the dephosphorylation process: in immature neurons (DIV7), NMDA treatment promoted a small increase in GluN2B-pS1480, rather than the expected decrease that we observed beginning at DIV14, and became increasingly enhanced throughout neuronal maturation (Figure 3H).

### GluN2B-pS1480 dephosphorylation is mediated by a population of PP1 not activated during LTD

Because GluN2B-pS1480 receptors are exclusively expressed at extrasynaptic membranes (Figure 1B) and there is no translocation of PP1 to the surface upon NMDA treatment (Figure 3D/E), we hypothesized that the PP1 population mediating this reaction is also extrasynaptically located. This idea would be consistent with the need of the activation of extrasynaptic NMDARs to induce dephosphorylation (Figure 2C) and it would explain the differential outcome of NMDA treatment in immature cultures (Figure 3H), as the compartmentalization between synaptic and extrasynaptic sites has not yet occurred. We tested this possibility by isolating GluN2B-containing protein complexes from adult rat brain extrasynaptic membranes and performing co-immunoprecipitation experiments with an anti-GluN2B antibody. As shown in Figure 3I, both PP1 and I-2 co-immunoprecipitated with GluN2B, demonstrating that they are part of the same extrasynaptic protein complex. Whereas GluN2B was enriched ~2-fold in our co-IP experiments, the enrichment of PP1 was higher (~4-fold) (Figure 3I). This indicates the existence of multiple copies of PP1 associated with a single GluN2B, likely indirectly, through the binding to the several “target specifiers” that link PP1 with GluN2B, including proteins like spinophilin (Baucum et al., 2013), and yotiao (Lin et al., 1998). Together, our results indicate that extrasynaptic GluN2B-containing NMDARs interact with PP1 and I-2, likely with the involvement of extrasynaptic PP1 target specifiers.

The existence of an extrasynaptic pool of PP1 (Fig 3I) in combination with our data showing the lack of PP1 recruitment to synapses after NMDA treatment (Figures 3D/E) is in sharp contrast with the reported synaptic recruitment of PP1 after the induction of Long-term Depression (LTD) (Morishita et al., 2001), suggesting the existence of distinct PP1 populations that can be differentially controlled. To test this possibility, we activated synaptic PP1 (Hu et al., 2007) by inducing chemical LTD (cLTD) in ex vivo hippocampal slices from adult (>P70) mice (Babiec et al., 2014). As a control, we quantified the levels of GluA1 phosphorylation at S845 (GluA1-pS845), as compelling evidence has demonstrated synaptic PP1-mediated GluA1-pS845 dephosphorylation (Diering and Huganir, 2018). We also quantified the levels of GluN2B-pS1480 in the same samples and, as shown in Figure 3J, we found a drastic difference between the regulation of the PP1-mediated GluN2B and GluA1 dephosphorylation after activation of synaptic PP1: whereas levels of GluA1-p845 decreased as expected, the levels of GluN2B-pS1480 increased after cLTD, presumably due to activation of synaptic NMDARs.

Together with our developmental data, these findings suggest that the GluN2B-pS1480 dephosphorylation reaction is mediated by the strict spatiotemporal trafficking of PP1 to extrasynaptic membranes.

### PP1 activity modulates NMDAR synaptic localization

If our hypothesis that GluN2B-pS1480 maintains NMDARs at extrasynaptic membranes is correct, then modulation of PP1 activity against GluN2B S1480 should cause changes in both synaptic signaling and synaptic GluN2B receptor content (Figure 4A). To test this idea, we first examined changes in ERK phosphorylation (pERK), because compelling evidence demonstrates that activation of synaptic NMDAR leads to an increase in pERK, whereas extrasynaptic NMDAR activation decreases pERK levels (Hardingham and Bading, 2010). Specifically, we modulated GluN2B-pS1480 in cultured neurons with NMDA (50 µM for 10 min) or OA (50 nM for 45 min) to induce the potential redistribution of NMDARs and, after drug withdrawal, we increased synaptic activity in the culture using a standard pharmacological approach [20 µM bicuculline and 100 µM 4-AP incubation (Bic/4-AP)]. As expected, cells treated with Bic/4-AP showed a robust increase in ERK phosphorylation that, consistent with our hypothesis, was potentiated in neurons pre-treated with NMDA. Conversely, cells pre-treated with OA to block PP1 activity displayed a reduced increase in ERK phosphorylation after Bic/4-AP treatment in comparison with control cells (please, note the expected elevated basal ERK phosphorylation in OA pre-treated neurons in comparison with controls) (Figure 4B).

**FIGURE 4:**
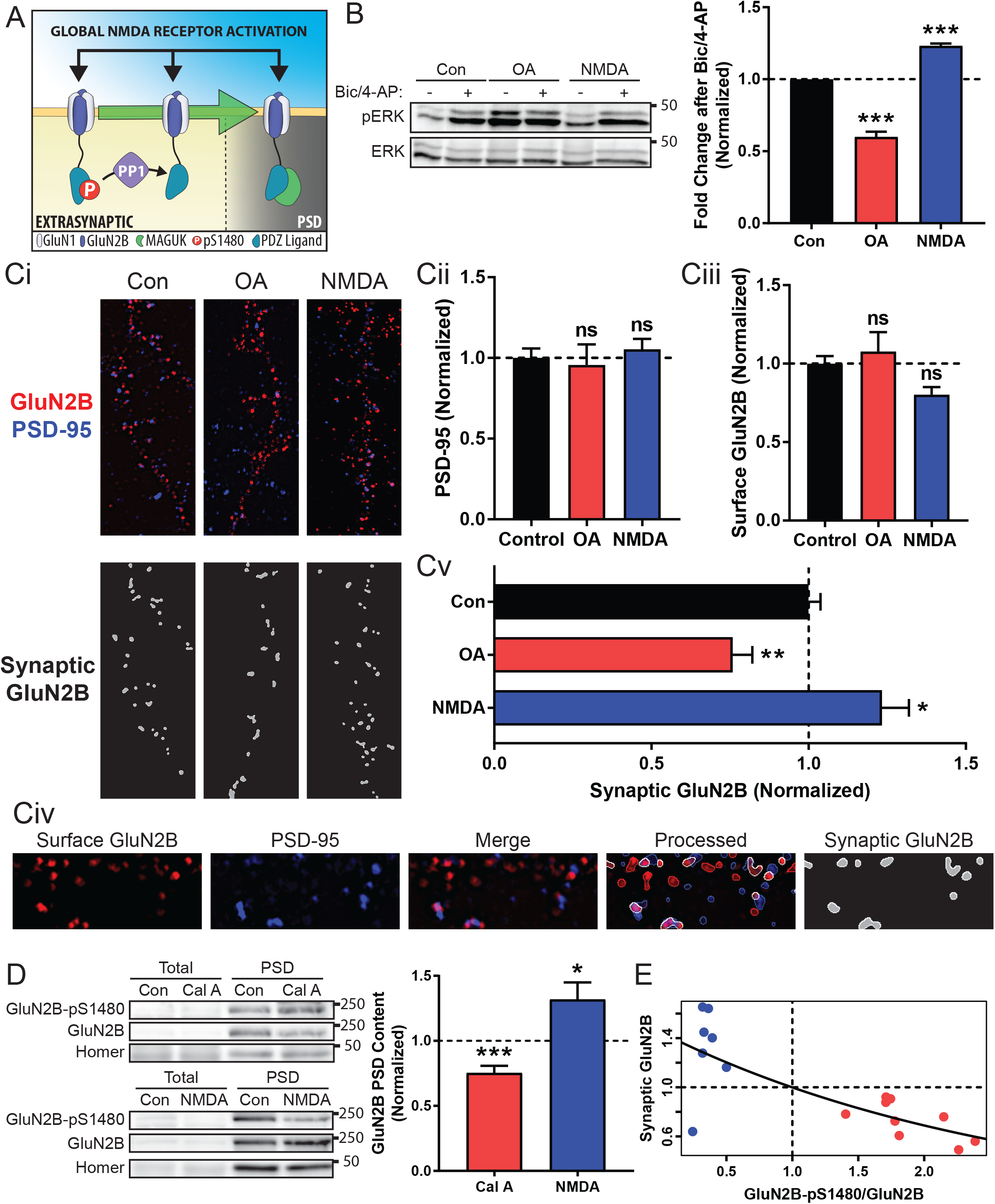
PP1 activity modulates NMDAR synaptic localization. All graphs represent mean ± SEM. *p<.05, **p<.01, ***p<.001 in a Mann-Whitney U test. **A)** Global synaptic and extrasynaptic NMDAR activation triggers PP1 activity to promote dephosphorylation of GluN2B which modulates its synaptic content. **B)** DIV18-21 cortical neurons were pre-treated with OA (50 nM for 45 min) or NMDA (50 µM for 10 min) and synaptic NMDAR stimulation performed by incubating with 20 μM Bicuculline and 100 μM 4-AP for 30 min. Levels of pERK and ERK were evaluated by immunoblotting. N=7 **C)** Structured Illumination Microscopy (SIM) super-resolution micrographs were acquired in GluN2B WT transfected neurons (Ci). After labeling with proper antibodies, total number of PSD-95 (Cii) and surface-expressed GluN2B (Ciii) puncta was quantified. To analyze the synaptic expression of exogenous receptor, Nikon NIS-Elements software identified colocalized puncta (synaptic GluN2B, white) [containing surface GFP-GluN2B (red) and endogenous PSD-95 (blue)] (Cv). Values are presented synaptic GluN2B vs total GluN2B puncta after pharmacological manipulations (Cv). Samples from each processing step are presented in Civ. N=4, n (Con, OA, NMDA)=55, 18, 29 **D)** Postsynaptic Density (PSD) fractions were isolated from acute hippocampal slices from P70-96 mice treated with NMDA (50 µM for 15 min) or Cal A (100 nM for 30 min to 1 hour) to promote or inhibit GluN2B-pS1480 dephosphorylation. Levels of GluN2B-pS1480 and synaptic GluN2B were analyzed by immunoblotting and normalized to the synaptic marker homer. N (Cal A, NMDA)=9, 7 **E)** Model to predict synaptic GluN2B content based on GluN2B-pS1480 levels. Blue circles represent data from acute slices treated with NMDA (50 µM for 15 min) and red circles represent slices treated with Cal A (100 nM for 30 min to 1 hour). Data points were fitted to an exponential regression. Trendline follows the equation: 1.4478*e*^−.37118(GluN2B-pS1480/GluN2B)^ (p<.001).

We next employed super-resolution microscopy in primary hippocampal cultures after transfection of GFP-GluN2B to investigate its synaptic expression after pharmacological treatment. Specifically, we labeled overexpressed receptors located at the surface and endogenous PSD-95 (as a synaptic marker) and used Structured Illumination Microscopy (SIM) to acquire images of ~120 nm resolution (Figure 4Ci). We first quantified the number of PSD-95 puncta and found that the pharmacological manipulations do not affect the global number of synaptic sites (Figure 4Cii). Because we have shown that NMDA treatment increases the surface expression of GluN2B (surface/intracellular) (Figure S2), we also quantified the net number of GluN2B puncta expressed at the surface of the analyzed neurons to be sure that they display a comparable level of surface-expressed GluN2B and, therefore, we can utilize consistent analytical parameters in all the conditions (Figure 4Ciii). Finally, we generated a mask containing the overlapped areas between GluN2B and PSD-95 puncta to identify synaptic NMDARs (Figure Civ). Quantification of the overlapped GluN2B/PSD-95 area revealed that incubation with NMDA to promote GluN2B-pS1480 dephosphorylation increased the synaptic expression of GFP-GluN2B, whereas OA treatment, which enhances GluN2B-pS1480, decreased GluN2B synaptic content (Figure 4Cv).

To confirm these data in a more intact preparation, we employed *ex vivo* PP1 manipulation in acute hippocampal slices. Slices treated with NMDA induced dephosphorylation of GluN2B S1480. Concurrent with this change in phosphorylation, we observed a robust increase in the normalized synaptic content of endogenous GluN2B. Conversely, slices treated with calyculin A, the most effective PP1 inhibitor in acute slices, displayed an increase in GluN2B-pS1480 and a concomitant decrease in GluN2B synaptic content (Figure 4D). To unbiasedly assess the importance of GluN2B-pS1480 in determining GluN2B synaptic localization, we first tested the strength of correlation between normalized GluN2B-pS1480 levels and normalized GluN2B synaptic content using the data obtained with pharmacologically-treated brain slices (Figure 4D). Specifically, we calculated the Spearman’s rank correlation and found a strong association between these parameters (p<.01). We, therefore, fit the data to an exponential model with the following equation: Synaptic GluN2B = 1.4478*e*^−.37118(GluN2B-pS1480/GluN2B)^ (p<.001), which predicts synaptic NMDAR content based on levels of GluN2B-pS1480 (Figure 4E). Together, these data demonstrate that PP1 activity against the GluN2B PDZ-ligand is a crucial regulator of GluN2B-containing NMDAR trafficking and controls NMDAR synaptic content in mature neurons.

## DISCUSSION

In this study, we identify the PP1-mediated dephosphorylation of the PDZ-ligand of GluN2B (at S1480) as a critical determinant for controlling NMDAR synaptic content in mature neurons. Specifically, we show that phosphorylated NMDARs are located at extrasynaptic membranes and complex with PP1. Global activation of NMDARs results in PP1 activation and dephosphorylation of GluN2B. Our data support a model in which dephosphorylated receptors are integrated into the PSD to modulate NMDAR synaptic content. Our observations indicate that insertion of NMDAR into PSDs requires the regulated dephosphorylation of GluN2B-pS1480. This mechanism is consistent with the observation that GluN2B-containing, but not GluN2A-contaning, NMDARs are subject to lateral diffusion between synaptic and extrasynaptic sites (Dupuis et al., 2014), as we have previously demonstrated that GluN2A PDZ-ligand is not efficiently phosphorylated by CK2 (Sanz-Clemente et al., 2010).

Our findings expand our previous work characterizing the molecular events controlling the synaptic activity-dependent phosphorylation of the PDZ-ligand of GluN2B (Lussier et al., 2015). Briefly, we showed that casein kinase 2 (CK2) phosphorylates GluN2B S1480 to disrupt the interaction of NMDARs with MAGUK proteins and promote NMDAR synaptic clearance and internalization (Chung et al., 2004; Sanz-Clemente et al., 2013a; Sanz-Clemente et al., 2010). We have now shown that NMDARs phosphorylated at S1480 are recycled to plasma membrane and identified the mechanism underlying the insertion of receptors to the PSD. Together, our data show that the dynamic regulation of the phosphorylation state of the GluN2B PDZ-ligand is a key modulator of the activity-dependent modifications in the levels of synaptic NMDARs (Dore et al., 2017; Hunt and Castillo, 2012). Given the essential functional and regulatory roles displayed by synaptic NMDARs, modification of synaptic NMDAR density has emerged as a key mechanism for controlling neuronal function. The most evident consequence of this process is the modulation of metaplasticity by modifying the induction threshold of AMPAR-plasticity. However, other neuronal properties, such as the integration of synaptic inputs or the control of homeostatic plasticity may also be affected (Dore et al., 2017; Hunt and Castillo, 2012). The precise molecular mechanisms underlying the modulation of NMDAR synaptic content remain unclear, although the involvement of the co-activation of NMDARs and GPCRs (i.e., mGluR1/5 or mAChRs) have been reported (Dore et al., 2017). Here, we identify a molecular mechanism that controls the synaptic recruitment of extrasynaptic NMDARs to synaptic sites, a process previously identified to contribute to increase NMDAR synaptic content (Dupuis et al., 2014; Harney et al., 2008). Regulation of synaptic/extrasynaptic NMDAR exchange is important because elevated extrasynaptic NMDAR signaling triggers excitotoxicity and alterations in the NMDAR balance has been associated with several neurological disorders including Huntington’s disease, traumatic brain injury (TBI), and epilepsy (Parsons and Raymond, 2014). Therefore, impairments in the PP1-mediated regulation of NMDAR trafficking may contribute to the pathophysiology of these disorders.

PP1 is a ubiquitously expressed serine/threonine phosphatase, although in neurons, it is enriched in dendritic spines (Strack et al., 1999). Our finding that activation of PP1 increases NMDAR synaptic content by dephosphorylating GluN2B PDZ-ligand is unexpected as phosphatases, and particularly PP1, have been associated with the depression of synaptic activity rather than its potentiation (Woolfrey and Dell’Acqua, 2015). In fact, compelling evidence supports a model in which PP1 induces AMPAR-LTD (reduction of synaptic AMPAR content) by dephosphorylating the GluA1 subunit of AMPAR (at S845) and CaMKII (at T256) (Siddoway et al., 2014). Although it can occur independently, modification in NMDAR and AMPAR synaptic content are often co-induced (Hunt et al., 2013). How, then, can PP1 activity drive opposite effects in synaptic NMDARs and AMPARs density (i.e., increasing NMDAR but decreasing AMPAR synaptic content)? We propose that the activation of different pools of PP1 is a critical factor for the different outcomes. Therefore, the stimuli classically used to induce AMPAR-LTD would activate synaptic resident-PP1, but not the PP1 pool associated with GluN2B at the extrasynaptic membranes (as extrasynaptic NMDARs would not be activated). Conversely, strong stimuli required for the increase of synaptic NMDAR content would cause a robust release of glutamate, enough to spillover and activate both synaptic and extrasynaptic NMDARs. This would result in the increase in both AMPAR (by CaMKII activation and AMPAR phosphorylation) and NMDAR (by PP1 activation and dephosphorylation of GluN2B-pS1480) synaptic content. The existence of different pools of PP1 that can be differentially activated is consistent with the differential regulation of the PP1-mediated dephosphorylation of GluN2B-pS1480 and GluA1-pS845 (Figure 3J) and is supported by the fact the stimulus used to induce AMPA-LTD in culture recruits PP1 to the synaptic sites (Morishita et al., 2001), whereas our biochemical and TIRF data indicate that global NMDAR activation does not (Figures 3D/E). In summary, our data reveal a critical role for PP1 in controlling NMDAR synaptic content in mature neurons by mediating the dephosphorylation of GluN2B upon global activation of NMDARs.

## Acknowledgments

We thank Alec M. Chiu for assistance with statistical analysis and Luca Zangrandi for technical assistance. We also thank the Northwestern Center for Advanced Microscopy for their assistance in planning and analyzing imaging experiments. This research was supported by NIGMS (T32GM008061) (A.M.C.), NIMH (K08MH100562) (J.A.G.) and NIA (K99AG041225) and a NARSAD Young Investigator Grant from the Brain & Behavior Research Foundation (#24133) (A.S.-C.).

## CONTACT FOR REAGENT AND RESOURCE SHARING

Further information and requests for resources and reagents should be directed to and will be fulfilled by the Lead Contact, Antonio Sanz-Clemente

## EXPERIMENTAL MODEL AND SUBJECT DETAILS

The care and use of animals were in accordance to the guidelines set by the Northwestern University IACUC and UCSF Animal Research Advisory Committee.

### Primary neuronal cultures

Primary neuronal cultures were generated from E18 Sprague-Dawley rats. Dissociated neurons were plated on poly-D-lysine coated plates or coverglasses Cells were maintained in Neurobasal media (Gibco) supplemented with B-27 (Gibco) and glutamine (20 mM) (Gibco). For biochemical experiments, primary cortical neuronal cultures at a density of ~150,000 cell/cm^2^ were used due to the ease of generating enough material for analysis. For imaging-based experiments, hippocampal primary neuronal cultures were utilized at a density of ~40,000 cell/cm^2^.

### Animals

C57/B6 male and female mice were mainlined by the Northwestern Center for Comparative Medicine. Mice were group housed (up to 5 mice per cage), housed on a 14/10 light-dark schedule, and had access to food and water *ad libitum*. Mice of the indicated ages were used to generate either slice culture or acute slices for ex vivo experiments.

## METHOD DETAILS

### Lentivirus Generation

Lentiviruses were generated by transfecting Lenti-X cells (Clontech) with psPAX2, pMD2.G, and the pLVX backbone containing the indicated construct. The viral supernatant was harvested 48 and 72 hours post transfection and concentrated using polyethylene glycol for 24 hours. Concentrated virus was resuspended 100x in PBS and stored at −80ºC until use.

### Immunoflouresence Microscopy

For recycling experiments, hippocampal neurons were transfected at DIV7 with GFP-tagged GluN2B (WT or mutants), and surface-expressed receptors were labeled with anti-GFP antibody for 15 min at RT 4-7 days post transfection. Following internalization receptors (30 min at 37°C), remaining surface receptors were blocked with Fab (20 µg/ml) for 20 min at RT. After allowing for recycling of receptors back to the surface (45 min at 37°C), cells were washed and fixed with 4% PFA in PBS containing 4% sucrose. Surface receptors were labeled with Alexa 555-conjugated secondary antibody (shown in white for clarity). The intracellular pool of receptors was identified by permeabilizing cells with 0.25% Triton X-100 and labeling anti-GFP tagged receptors with Alexa 633-conjugated secondary antibodies (shown in green). Cells were imaged on a Zeiss LSM 510 confocal microscope. Serial optical sections collected at 0.35 μm intervals were used to create maximum projection images. Quantification was performed by analyzing the fluorescence intensity of 3-4 independent areas per neuron using MetaMorph 6.0 software (Universal Imaging Corp) and is presented as ratio surface/intracellular intensities (mean ±SEM).

For surface expression experiments, cells were transfected on DIV14 and subject to pharmacologic manipulation immediately prior to labeling with an anti-GFP antibody for 15 min at RT surface receptors and subsequent labeling with Alexa 555-conjugated secondary antibody. The intracellular pool of receptors was identified by permeabilizing cells with 0.25% Triton X-100 and labeling anti-GFP tagged receptors with Alexa 647-conjugated secondary antibodies. Images were acquired on a Nikon A1 confocal microscope. Serial optical sections collected at 0.35 μm intervals were used to create maximum projection images. Quantification was performed by analyzing the fluorescence intensity of 5 independent areas per neuron using FIJI and is presented as ratio surface/intracellular intensities (mean ±SEM).

### Electrophysiology in Organotypic Slice Cultures

Double-floxed GluN2A/GluN2B (*Grin2a*^fl/fl^*Grin2b*^fl/fl^) mice were generated as previously described (Akashi et al., 2009; Gray et al., 2011; Mishina and Sakimura, 2007). Cultured slices were prepared and transfected as previously described (Schnell et al., 2002). Briefly, hippocampi were dissected from P7 *Grin2a*^fl/fl^*Grin2b*^fl/fl^ mice and biolistically co-transfected after 2-4 days in culture with pFUGW-Cre:mCherry (expressing a nuclear targeted Cre:mCherry fusion protein) and either pCAGGS-GFP or pCAGGS-GluN2B-S1480E-IRES-GFP. Slices were cultured for an additional 14–20 days before recording. Slices were recorded in a submersion chamber on an upright Olympus microscope, perfused in room temperature normal ACSF saturated with 95% O2/5% CO2. Picrotoxin (0.1 mM) and NBQX (10 μM) were added to the ACSF to block GABAA and AMPA receptors respectively. CA1 pyramidal cells were visualized by infrared differential interference contrast microscopy and transfected neurons were identified by epifluorescence microscopy. The intracellular solution contained (in mM): CsMeSO_4_ 135, NaCl 8, HEPES 10, Na- GTP 0.3, Mg-ATP 4, EGTA 0.3, and QX-314 5. Cells were recorded with 3 to 5MΩ borosilicate glass pipettes, following stimulation of Schaffer collaterals with bipolar electrodes placed in stratum radiatum of the CA1 region. Series resistance was monitored and not compensated, and cells in which series resistance varied by 25% during a recording session were discarded. Synaptic responses were collected with a Multiclamp 700B amplifier (Axon Instruments, Foster City, CA), filtered at 2 kHz, digitized at 10 Hz. All paired recordings involved simultaneous whole-cell recordings from transfected neuron and a neighboring untransfected neuron. NMDAR-EPSCs were recorded at +40 mV in the presence of 10 μM NBQX. The stimulus was adjusted to evoke a measurable, monosynaptic EPSC in both cells. Whole cell NMDA responses were evoked with 100 µM NMDA (in ACSF) pressure-ejected (puffed) from a glass pipette by a Picospritzer III (Parker-Hannifin, Hollis, NH). Slices were aligned such that NMDA was puffed over the soma of the cell pair and perfused along the apical dendrites following the bath flow direction. Paired amplitude data were analyzed with a Wilcoxon signed-rank test and comparison of paired data groups were performed using a Mann-Whitney U test. Linear regressions were obtained using the least-squares method. All errors bars represent standard error measurement (SEM).

### Generation of Neuronal Membrane Fractions

The crude membrane fraction was generated as previously described (Sanz-Clemente, et. al. 2010). Briefly, cortical neurons were harvested in PBS containing 0.1 mM CaCl_2_ and 1 mM MgCl_2_, and pelleted by centrifugation at 20,000 g. Pellets were resuspended in hypotonic buffer (10 mM Tris, 1 mM NaVO_4_) and sonicated. Membranes were isolated by centrifugation at 20,000 g and resuspended in SDS loading buffer.

### Biochemical Subcellular Fractionation

Biochemical subcellular fraction followed established protocols. Briefly, brain tissue or cells were homogenized in homogenization buffer (0.32M HEPES-buffered sucrose) containing protease (ROCHE) and phosphatase (Sigma) inhibitors and centrifuged for 10 min at 1,000 g to remove nuclei and large debris. The supernatant was centrifuged at 10,000 g for 15 min to generate the synaptosomal fraction (P2). P2 was resuspended in hypotonic buffer, sonicated, and centrifuged at 25,000 g to generate the SPM fraction (P3). To separate PSDs from the extrasynaptic fraction, SPMs were resuspended in PBS containing 1% Triton X-100 and ultracentrifuged at 100,000 g for 1 hour. Pellets containing the PSD were resuspended in PBS.

### Total Internal Reflection Microscopy

For total internal reflection microscopy, primary hippocampal neurons were plated in glass bottom dishes (Matek). On DIV 10, cells were transfected with either GFP-PP1α/γ1 or GFP-CaMKII. Cells were imaged on DIV 14 on a NIKON X1 spinning disk confocal microscope. During imaging, cells were maintained at 37°C with appropriate humidity and gas using a Tokai Hit incubation chamber. A timelapse series captured images every 30s to establish at least 5m of baseline and continued capturing images for 10 minutes following treatment with 50 μM NMDA. Timelapse series were acquired with NIKON elements software and changes in fluorescence at each timepoint were analyzed using FIJI. Changes in fluorescence intensity from baseline are reported as mean ± SEM.

### Co-immunoprecipitation from extrasynaptic membranes

For co-immoprecipitation, SPM fraction was obtained from rat brain cortex as explained before, dissolved in PBS 1% Triton X-100 and ultracentrifuged at 100,000 g for 1 hour. The pellet (PSD) was discarded, and the “extrasynaptic membranes” fraction (SN) was incubated with 2 µg anti-GluN2B antibody (or IgG as a control) for 1 hour at 4ºC with rotation. BSA-preblocked protein A-Sepharose beads were added to the lysate and incubated for 1 additional hour. After 3×10 minutes washes with cold PBS 1% TX-100, samples resuspended in SDS loading buffer.

### Immunoblotting

Follow preparation, samples were then run on SDS-PAGE and immunoblotted with the indicated antibodies. For phosphorylation-state specific and total immunoblots, the same membrane was used following antibody stripping. Quantification was performed by FIJI and presented as mean ± SEM.

The uncropped blots utilized for assembling the figures in this study can be found at: https://data.mendeley.com/datasets/769vrs9v2v/draft?a=7901a11c-4c68-4817-ad94-7b09aab4ab1a

### Superresolution Microscopy

For super-resolution microscopy, hippocampal primary cultured neurons were transfected with GFP-GluN2B at DIV14 and processed at DIV21-23. Surface-expressed receptors were labeled with an anti-GFP antibody and cells were fixed and permeabilized as before. Endogenous PSD-95 was labeled with a specific antibody as a synaptic marker. Super-resolution images were acquired using a Nikon N-SIM Structured Illumination Super-resolution Nikon-SIM microscope and reconstructed with NIS-Elements software. One secondary dendrite was selected and imaged from each neuron. Reconstruction parameters (0.74;0.83;0.21) were kept consistent throughout the experiment. Imaging and reconstruction parameters were empirically determined with the assistance of the expertise of the Nikon Imaging Center at Northwestern University. Reconstructed images were then opened in Fiji software. After thresholding both channels, a colocalized channel was created containing the overlapped area of GFP-GluN2B and PSD95 channels. Particle analysis were conducted on all three channels to get the total area and number of GFP-GluN2B, PSD95 and colocalized proteins independently. Synaptic GluN2B was determined by the ratio between colocalized puncta area and total GFP-GluN2B puncta area expressed on cell surface and are reported as mean ± SEM.

### Pharmacologic Manipulation of Acute Slices

Acute cortical slices were generated from adult (8 to 14 week) C57/B6 mice. Mice were anesthetized with isoflurane prior to decapitation. Coronal slices (300 µm thick) were cut in ice-cold artificial cerebrospinal fluid containing the following composition (in mM): NaCl 130, KCl 3.5, Glucose 10, NaH_2_PO_4_ 1.25, NaHCO_3_ 24, CaCl_2_ 1, MgCl_2_ 2. The slices were incubated for 30 min at 30-32°C and allowed to rest for at least 1 hour at room temperature in normal ACSF containing (in mM): NaCl 130, KCl 3.5, Glucose 10, NaH_2_PO_4_ 1.25, NaHCO_3_ 24, CaCl_2_ 2, MgCl_2_ 1. For pharmacologic manipulation, slices were incubated at 32°C and treated with either 50 μM NMDA for 15 minutes (NMDA), 100 nM Cal A for 30-60 min (Cal A), or 20 μM NMDA for 3 minutes with 30 minutes of recovery in drug-free ACSF (cLTD). Following pharmacologic manipulation, slices were flash frozen and subject to subsequent lysis or fractionation.

### Modeling of GluN2B Synaptic content

Data from acute slices treated with either NMDA or Cal A was pooled. A Spearman’s rank correlation was calculated to verify association between normalized GluN2B-pS1480 levels and normalized synaptic GluN2B content. Subsequently, the data was fit to an exponential model following the form *Y* = *e*^*ax*+*b*^. Model was obtained by regressing the natural log of synaptic GluN2B content against GluN2B-pS1480 levels using the *lm* function in R. F-test was used to calculate statistical significance of the model.

Our antibody against phosphorylation state-specific GluN2B S1480 (Ac-CGHVYEKLSSIE(pS)DV-OH) was generated by New England Peptide. Antibodies against GluN2B and PSD-95 were obtained from Neuromab. Transferrin, CaMKII, and PP1γ were obtained from Thermo Fisher. PP1 antibody was obtained from Santa Cruz. All drugs and inhibitors were obtained from Tocris Biosciences.

## QUANTIFICATION AND STATISTICAL ANALYSIS

Quantification methods for individual experiments are described as above in method details. Statistical tests and significance values are reported in figure legends. All data is presented as mean ± SEM. N represents number of experimental repeats. For imaging experiments, n represents the number of neurons sampled. Prism 6 (Graphpad) was used for all statistical tests. R was used to generate GluN2B mathematical model. Tests utilized include Mann-Whitney U Test, Kruskal-Wallis H Test, and F test. For all analysis, ns=non-significant, *p<.05, **p<.01, ***p<.001, ****p<.0001.

---

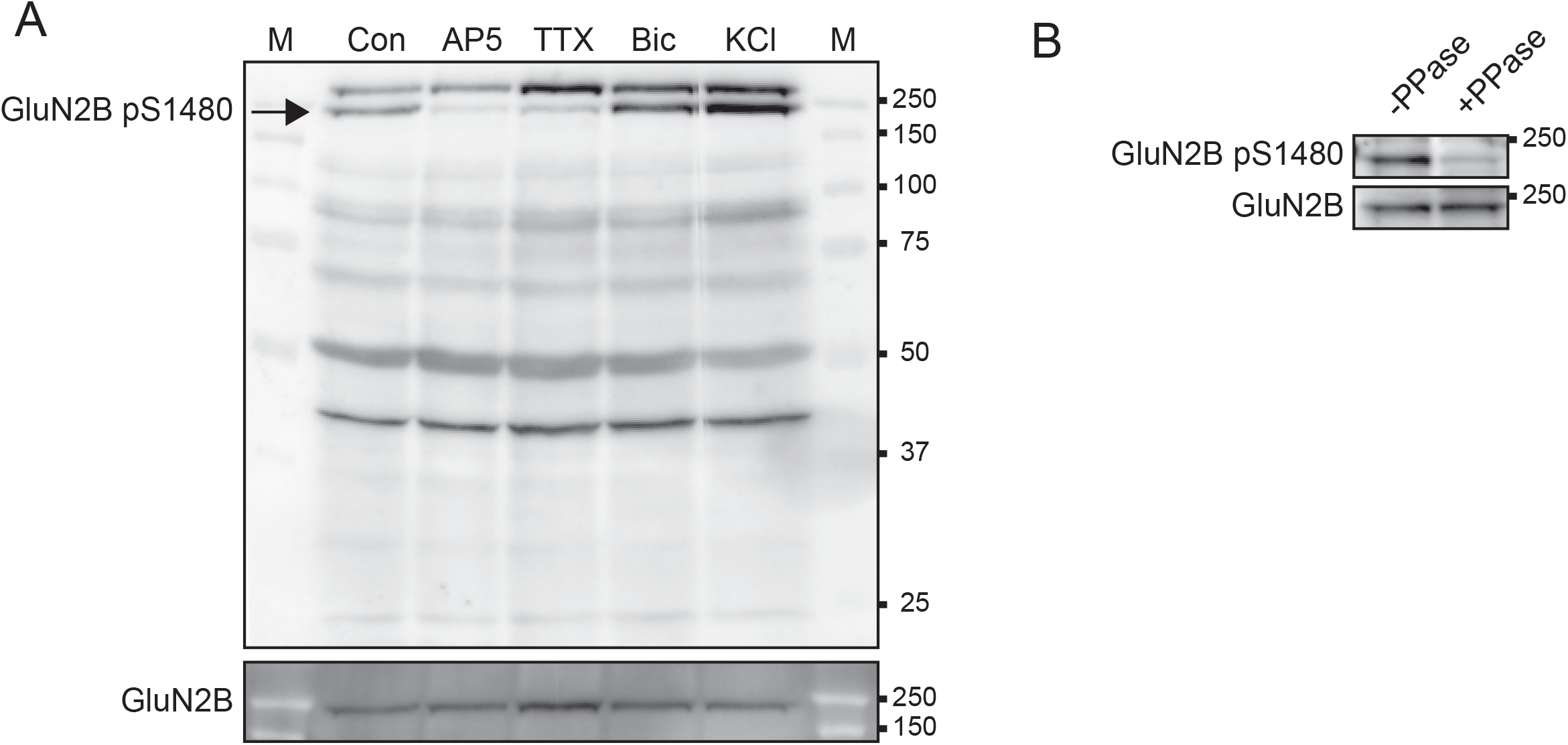

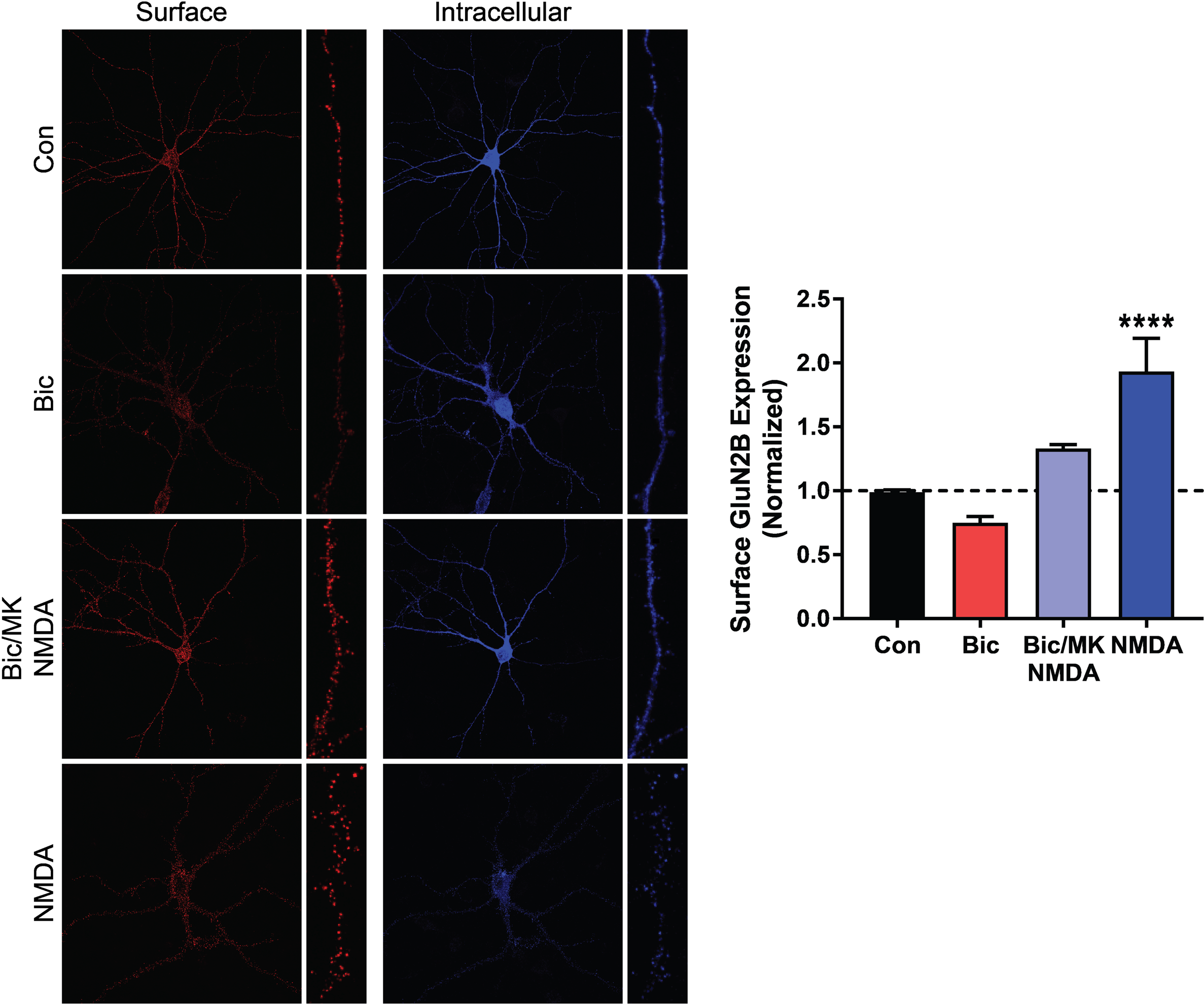

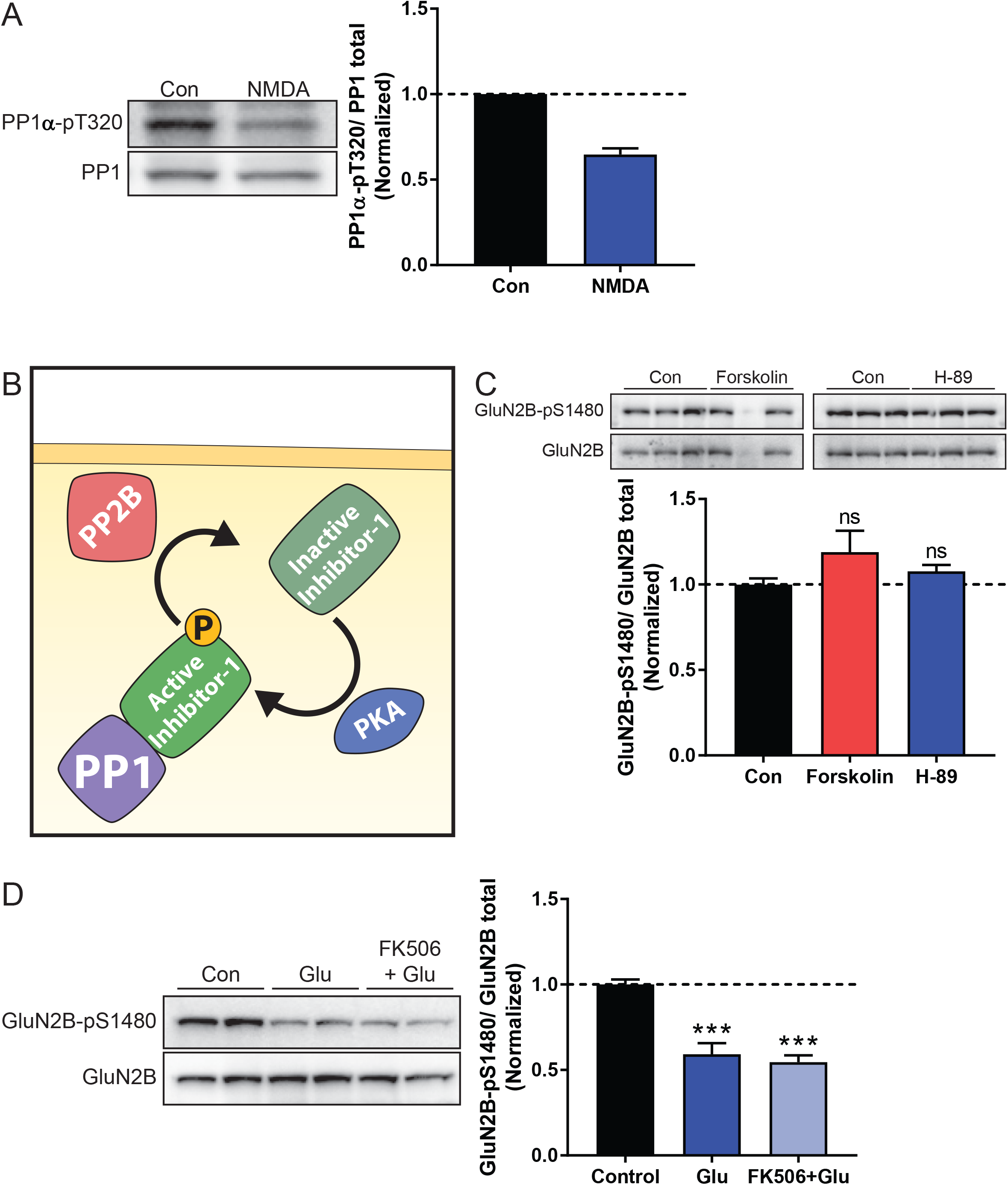

## REFERENCES

Babiec, W.E., Guglietta, R., Jami, S.A., Morishita, W., Malenka, R.C., and O’Dell, T.J. (2014). Ionotropic NMDA receptor signaling is required for the induction of long-term depression in the mouse hippocampal CA1 region. The Journal of neuroscience : the official journal of the Society for Neuroscience 34, 5285–5290.

Baucum, A.J., 2nd, Brown, A.M., and Colbran, R.J. (2013). Differential association of postsynaptic signaling protein complexes in striatum and hippocampus. Journal of neurochemistry 124, 490–501.

Chen, B.S., Gray, J.A., Sanz-Clemente, A., Wei, Z., Thomas, E.V., Nicoll, R.A., and Roche, K.W. (2012). SAP102 mediates synaptic clearance of NMDA receptors. Cell reports 2, 1120–1128.

Chung, H.J., Huang, Y.H., Lau, L.F., and Huganir, R.L. (2004). Regulation of the NMDA receptor complex and trafficking by activity-dependent phosphorylation of the NR2B subunit PDZ ligand. The Journal of neuroscience : the official journal of the Society for Neuroscience 24, 10248–10259.

Cohen, P.T. (2002). Protein phosphatase 1––targeted in many directions. J Cell Sci 115, 241–256.

Diering, G.H., and Huganir, R.L. (2018). The AMPA Receptor Code of Synaptic Plasticity. Neuron 100, 314–329.

Dore, K., Stein, I.S., Brock, J.A., Castillo, P.E., Zito, K., and Sjostrom, P.J. (2017). Unconventional NMDA Receptor Signaling. The Journal of neuroscience : the official journal of the Society for Neuroscience 37, 10800–10807.

Dupuis, J.P., Ladepeche, L., Seth, H., Bard, L., Varela, J., Mikasova, L., Bouchet, D., Rogemond, V., Honnorat, J., Hanse, E., et al. (2014). Surface dynamics of GluN2B-NMDA receptors controls plasticity of maturing glutamate synapses. The EMBO journal 33, 842–861.

Eto, M., and Brautigan, D.L. (2012). Endogenous inhibitor proteins that connect Ser/Thr kinases and phosphatases in cell signaling. IUBMB Life 64, 732–739.

Glausier, J.R., Maddox, M., Hemmings, H.C., Jr., Nairn, A.C., Greengard, P., and Muly, E.C. (2010). Localization of dopamine- and cAMP-regulated phosphoprotein-32 and inhibitor-1 in area 9 of Macaca mulatta prefrontal cortex. Neuroscience 167, 428–438.

Hardingham, G.E., and Bading, H. (2010). Synaptic versus extrasynaptic NMDA receptor signalling: implications for neurodegenerative disorders. Nature reviews Neuroscience 11, 682–696.

Hardingham, G.E., Fukunaga, Y., and Bading, H. (2002). Extrasynaptic NMDARs oppose synaptic NMDARs by triggering CREB shut-off and cell death pathways. Nat Neurosci 5, 405–414.

Harney, S.C., Jane, D.E., and Anwyl, R. (2008). Extrasynaptic NR2D-containing NMDARs are recruited to the synapse during LTP of NMDAR-EPSCs. The Journal of neuroscience : the official journal of the Society for Neuroscience 28, 11685–11694.

Hendrickx, A., Beullens, M., Ceulemans, H., Den Abt, T., Van Eynde, A., Nicolaescu, E., Lesage, B., and Bollen, M. (2009). Docking motif-guided mapping of the interactome of protein phosphatase-1. Chem Biol 16, 365–371.

Hou, H., Sun, L., Siddoway, B.A., Petralia, R.S., Yang, H., Gu, H., Nairn, A.C., and Xia, H. (2013). Synaptic NMDA receptor stimulation activates PP1 by inhibiting its phosphorylation by Cdk5. The Journal of cell biology 203, 521–535.

Hu, X.D., Huang, Q., Yang, X., and Xia, H. (2007). Differential regulation of AMPA receptor trafficking by neurabin-targeted synaptic protein phosphatase-1 in synaptic transmission and long-term depression in hippocampus. The Journal of neuroscience : the official journal of the Society for Neuroscience 27, 4674–4686.

Hunt, D.L., and Castillo, P.E. (2012). Synaptic plasticity of NMDA receptors: mechanisms and functional implications. Current opinion in neurobiology 22, 496–508.

Hunt, D.L., Puente, N., Grandes, P., and Castillo, P.E. (2013). Bidirectional NMDA receptor plasticity controls CA3 output and heterosynaptic metaplasticity. Nature neuroscience 16, 1049–1059.

Ishihara, H., Martin, B.L., Brautigan, D.L., Karaki, H., Ozaki, H., Kato, Y., Fusetani, N., Watabe, S., Hashimoto, K., Uemura, D., et al. (1989). Calyculin A and okadaic acid: inhibitors of protein phosphatase activity. Biochem Biophys Res Commun 159, 871–877.

Lavezzari, G., McCallum, J., Dewey, C.M., and Roche, K.W. (2004). Subunit-specific regulation of NMDA receptor endocytosis. The Journal of neuroscience : the official journal of the Society for Neuroscience 24, 6383–6391.

Lin, J.W., Wyszynski, M., Madhavan, R., Sealock, R., Kim, J.U., and Sheng, M. (1998). Yotiao, a novel protein of neuromuscular junction and brain that interacts with specific splice variants of NMDA receptor subunit NR1. The Journal of neuroscience : the official journal of the Society for Neuroscience 18, 2017–2027.

Lussier, M.P., Sanz-Clemente, A., and Roche, K.W. (2015). Dynamic Regulation of N-Methyl-d-aspartate (NMDA) and alpha-Amino-3-hydroxy-5-methyl-4-isoxazolepropionic Acid (AMPA) Receptors by Posttranslational Modifications. The Journal of biological chemistry 290, 28596–28603.

Merrill, M.A., Chen, Y., Strack, S., and Hell, J.W. (2005). Activity-driven postsynaptic translocation of CaMKII. Trends in pharmacological sciences 26, 645–653.

Morishita, W., Connor, J.H., Xia, H., Quinlan, E.M., Shenolikar, S., and Malenka, R.C. (2001). Regulation of synaptic strength by protein phosphatase 1. Neuron 32, 1133–1148.

Paoletti, P., Bellone, C., and Zhou, Q. (2013). NMDA receptor subunit diversity: impact on receptor properties, synaptic plasticity and disease. Nature reviews Neuroscience 14, 383–400.

Papouin, T., and Oliet, S.H. (2014). Organization, control and function of extrasynaptic NMDA receptors. Philosophical transactions of the Royal Society of London Series B, Biological sciences 369, 20130601.

Parsons, M.P., and Raymond, L.A. (2014). Extrasynaptic NMDA receptor involvement in central nervous system disorders. Neuron 82, 279–293.

Sanz-Clemente, A., Gray, J.A., Ogilvie, K.A., Nicoll, R.A., and Roche, K.W. (2013a). Activated CaMKII couples GluN2B and casein kinase 2 to control synaptic NMDA receptors. Cell reports 3, 607–614.

Sanz-Clemente, A., Matta, J.A., Isaac, J.T., and Roche, K.W. (2010). Casein kinase 2 regulates the NR2 subunit composition of synaptic NMDA receptors. Neuron 67, 984–996.

Sanz-Clemente, A., Nicoll, R.A., and Roche, K.W. (2013b). Diversity in NMDA Receptor Composition: Many Regulators, Many Consequences. The Neuroscientist : a review journal bringing neurobiology, neurology and psychiatry 19, 62–75.

Siddoway, B., Hou, H., and Xia, H. (2014). Molecular mechanisms of homeostatic synaptic downscaling. Neuropharmacology 78, 38–44.

Siddoway, B.A., Altimimi, H.F., Hou, H., Petralia, R.S., Xu, B., Stellwagen, D., and Xia, H. (2013). An essential role for inhibitor-2 regulation of protein phosphatase-1 in synaptic scaling. The Journal of neuroscience : the official journal of the Society for Neuroscience 33, 11206–11211.

Strack, S., Kini, S., Ebner, F.F., Wadzinski, B.E., and Colbran, R.J. (1999). Differential cellular and subcellular localization of protein phosphatase 1 isoforms in brain. The Journal of comparative neurology 413, 373–384.

Szatmari, E., Habas, A., Yang, P., Zheng, J.J., Hagg, T., and Hetman, M. (2005). A positive feedback loop between glycogen synthase kinase 3beta and protein phosphatase 1 after stimulation of NR2B NMDA receptors in forebrain neurons. The Journal of biological chemistry 280, 37526–37535.

Tovar, K.R., and Westbrook, G.L. (2002). Mobile NMDA receptors at hippocampal synapses. Neuron 34, 255–264.

Virshup, D.M., and Shenolikar, S. (2009). From promiscuity to precision: protein phosphatases get a makeover. Mol Cell 33, 537–545.

Woolfrey, K.M., and Dell’Acqua, M.L. (2015). Coordination of Protein Phosphorylation and Dephosphorylation in Synaptic Plasticity. The Journal of biological chemistry 290, 28604–28612.

Zhang, J., Zhang, Z., Brew, K., and Lee, E.Y. (1996). Mutational analysis of the catalytic subunit of muscle protein phosphatase-1. Biochemistry 35, 6276–6282.

